# Accelerated clearing and molecular labeling of large tissue samples using magnetohydrodynamic force

**DOI:** 10.1101/819292

**Authors:** Joseph Dwyer, M. Desmond Ramirez, Paul S. Katz, Rolf O. Karlstrom, Joseph Bergan

**Affiliations:** Neuroscience and Behavior Graduate Program, University of Massachusetts Amherst; Department of Biology, University of Massachusetts Amherst; Department of Psychological and Brain Sciences, University of Massachusetts Amherst

## Abstract

Here we report a strategy to efficiently render opaque biological tissues transparent and demonstrate that this approach can be modified to rapidly label intact samples with antibodies for large volume fluorescence microscopy. This strategy applies a magnetohydrodynamic (MHD) force to accelerate the removal of lipids from and the introduction of antibodies into tissue samples as large as an intact adult mouse brain. This strategy complements a growing array of tools that enable high-resolution 3-dimensional anatomical analyses in intact tissues using fluorescence microscopy. MHD-accelerated clearing and MHD-accelerated antibody labeling are fast, reliable, inexpensive, and compatible with existing strategies for high-quality fluorescence microscopy of intact tissues.

## Introduction

Advances in microscopy make studying whole tissues with subcellular resolution possible, but generating suitable samples requires overcoming two major hurdles: opacity and infiltration. The presence of lipids in membranes causes light scattering, which makes it difficult or impossible to visualize structures deep in the tissue. Intact tissues also create a barrier for the delivery of agents that are needed to definitively label molecular features of interest. Here, we establish MHD force as an efficient mechanism to clear and label tissue samples for fluorescence microscopy.

Many biological samples, including the brain of most vertebrate animals, are recalcitrant to microscopy without first being cleared. Though chemical clearing has existed for over a century, these methods tended to quench fluorescence, making them unsuitable for high resolution fluorescence microscopy (Shultze, et al., 1897; Spalteholz, et al., 1914). More recent approaches to reduce opacity involve removing lipids from the tissue sample to reduce light scattering at lipid water interfaces (Chung, et al., 2013). When combined with genetically encoded fluorophores, these approaches have proven powerful for investigating anatomical relationships in a wide range of tissues and across a broad array of resolutions (Hama, et al., 2011; Lee, et al., 2016; Pan, et al., 2016; Susaki, et al., 2014; Susaki, et al., 2015; Murray, et al., 2015; Menegas, et al., 2015; Isogai, et al., 2014).

Immunohistochemistry is widely used because it is extremely effective at identifying molecules of interest in thinly sectioned biological samples. However, diffusion of antibodies into larger tissues (e.g. ∼1 cm^3^) takes inordinate amounts of time (weeks to months), depending on the size of the sample; even then, antibodies often cannot penetrate deep enough to label the sample completely and uniformly. Strategies have been developed to enhance the diffusion of antibodies into large tissue samples, however, these strategies can have serious drawbacks. CLARITY-based approaches provide high resolution imaging of endogenous fluorophores but can be difficult to implement with exogenous labels (e.g., IHC) in large samples (Chung, et al., 2013; Kim, et al., 2015; Lee, et al., 2016). Other approaches (e.g., iDISCO) use chemicals, microwaves, or electrophoresis to achieve optical transparency and allow fluorescence microscopy of large samples labeled with antibodies, but can quench endogenous fluorescent labels (Pan, et al., 2016; Renier, et al., 2014; Renier, et al., 2016; Susaki, et al., 2014; Hama, et al., 2015).

Here, we describe a strategy that addresses both major barriers to high resolution fluorescent imaging of large tissue samples. Magnetohydrodynamic (MHD) force, in combination with a conductive buffer and detergent, efficiently removes lipids from hydrogel-infused tissue and produces transparent samples excellent for fluorescence microscopy of genetically encoded fluorophores. The MHD force acts directly on ions inside the tissue sample, which simultaneously propels lipids out of the tissue and produces a constant flow of buffer through the tissue. This provides constant temperature regulation through the entire tissue. With this method, transparency of an intact mouse brain can be achieved within 2 days. MHD forces can subsequently be harnessed to drive antibodies into cleared tissues to bind to antigens. MHD-accelerated clearing and labeling works in both vertebrate (shown for mouse and zebrafish) and invertebrate (shown for the nudibranch mollusk *Berghia stephanieae*) species, providing a generalizable method to render intact tissue transparent and accelerate immunohistochemical labeling for fluorescence microscopy of intact tissues.

## Results

### Effects of MHD force

MHD force produces a linear increase in flow velocity that is not observed with the application of purely electrical force. To quantify the effects of the MHD force, we compared the movement of sodium alginate spheres in response to purely electrical or MHD forces. The MHD condition produced higher velocity flow over the electrical only condition for all tested non-zero voltages (Figure 1; p < 0.0001: t-test with Bonferroni correction). The difference between MHD and electric-only flow velocity increased as the applied voltage increased (Figure 1, Video 1).

**Figure 1:**
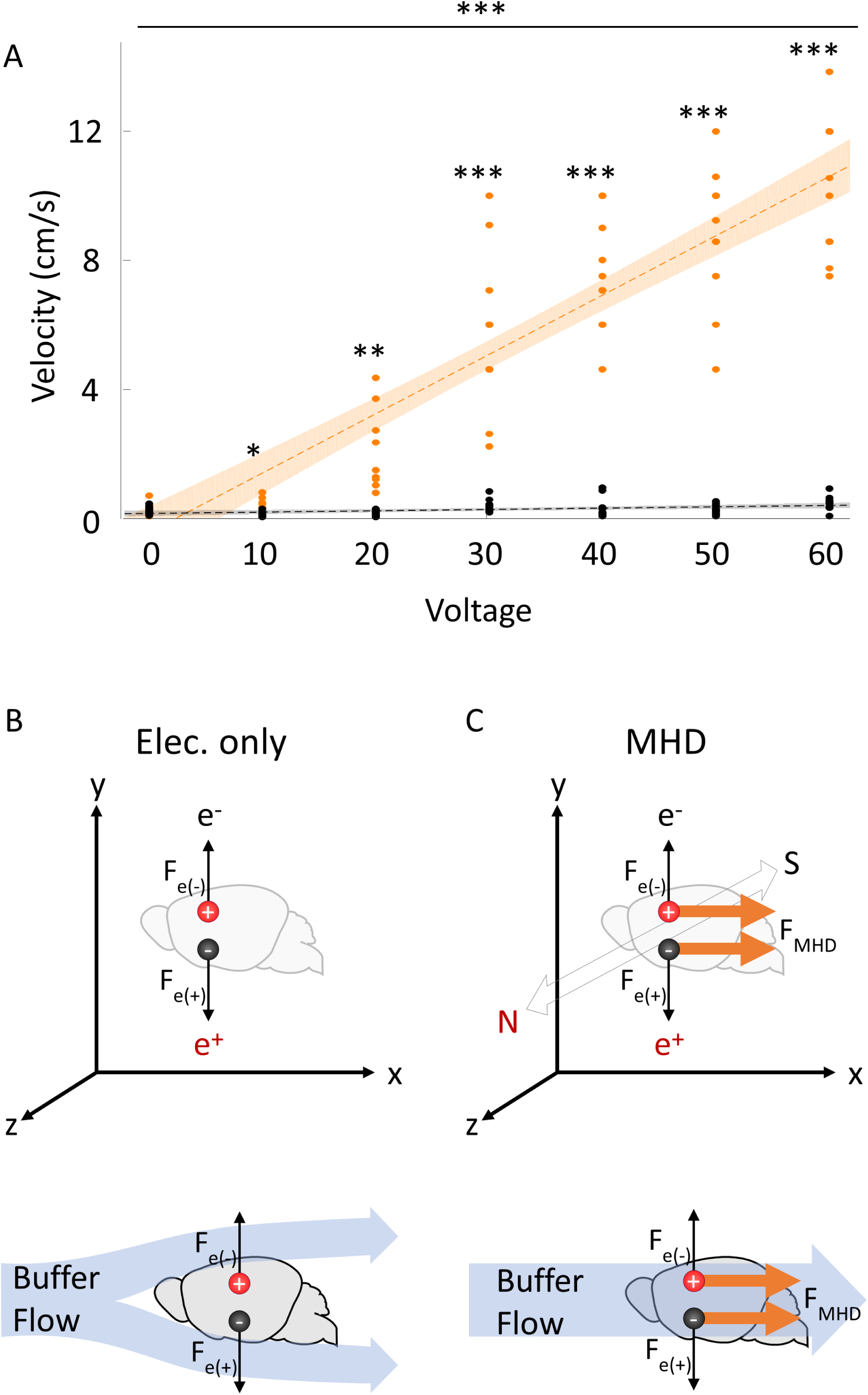
Comparison of voltage effects on buffer velocity between MHD and electrical forces. A) Velocity on sodium alginate spheres through the MHD-accelerated clearing device with (orange) and without a magnetic field (black; N = 7). MHD had a higher velocity than electric only all non-zero voltages (p < 0.0001; individual comparisons: *<0.01, **<0.001, ***<0.0001). A linear fit (y=mx+b) to each dataset is shown (shading 95% CI for the fitted line). B) Illustration of the forces produced by an electric field on cations (red) and anions (black). C) Illustration of the forces produced by MHD on cations and anions. Note that the MHD force is in the same direction for both cations and anions. Because the force on anions and cations is in a single direction, MHD produces bulk flow in the buffer and inside the tissue, whereas, an external pump is required when only using an electrical force.

### Tissue Clearing/Delipidation

MHD-accelerated clearing rendered a whole adult mouse brain transparent in as few as 15 hours (Figure 2). Pretreatment with passive incubation in SDS-containing clearing solution (37° C) for two days prior to MHD-accelerated clearing improved optical transparency in terms of both light transmission and effective clarity (Figure 2 – Figure supplement 1). Because pretreatment was gentler on tissue and reduced the time of MHD clearing required to achieve transparency, all subsequent tissue samples were passively cleared prior to active clearing (Figure 2 —Figure supplement 1).

**Figure 2:**
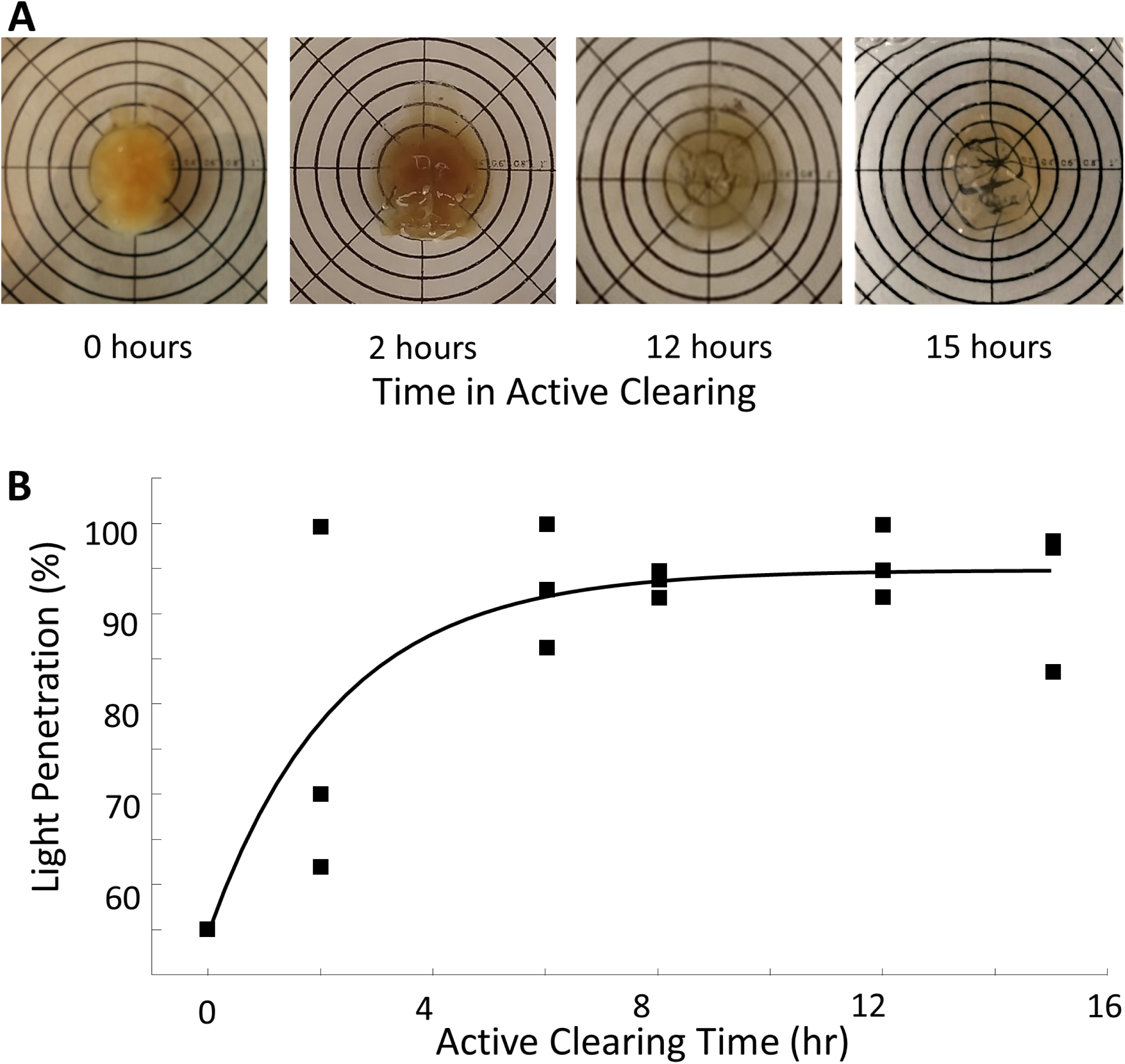
MHD-accelerated clearing of the intact mouse brain. A) Representative examples of intact cleared brains actively cleared with MHD for 0,2,12, and 15 hours and then equilibrated in RI-matching solution (N = 18). B) Measurements of the optical transparency of actively cleared mouse brains. Transparency was measured as percentage wide-spectrum light penetration through the tissue (curve fit with a saturating exponential).

MHD-accelerated clearing reliably rendered tissue samples optically transparent while also preserving genetically encoded fluorescent proteins (Figure 3). Of the 36 adult mouse brains used in this paper and the more than 50 cleared using this technique in multiple laboratories, all achieved transparency with no visible damage (Figure 2A). An intact adult mouse brain conditionally expressing GFP via EnvA-G-deleted rabies virus in aromatase expressing neurons was prepared using MHD-accelerated clearing and imaged on a Zeiss Z1 lightsheet microscope (Watabe-Uchida et al., 2016; Yao, et al., 2017). Sparse GFP cells were easily identified in the center of the brain (Figure 3B, C, D). Whole brain images resolved tissue architecture throughout the brain with subcellular resolution (Figure 4C, D). Higher magnification showed that fine processes, such as dendrites and axons, can be easily identified and analyzed several mm from surface of the brain (Figure 3D, Video 2).

**Figure 3:**
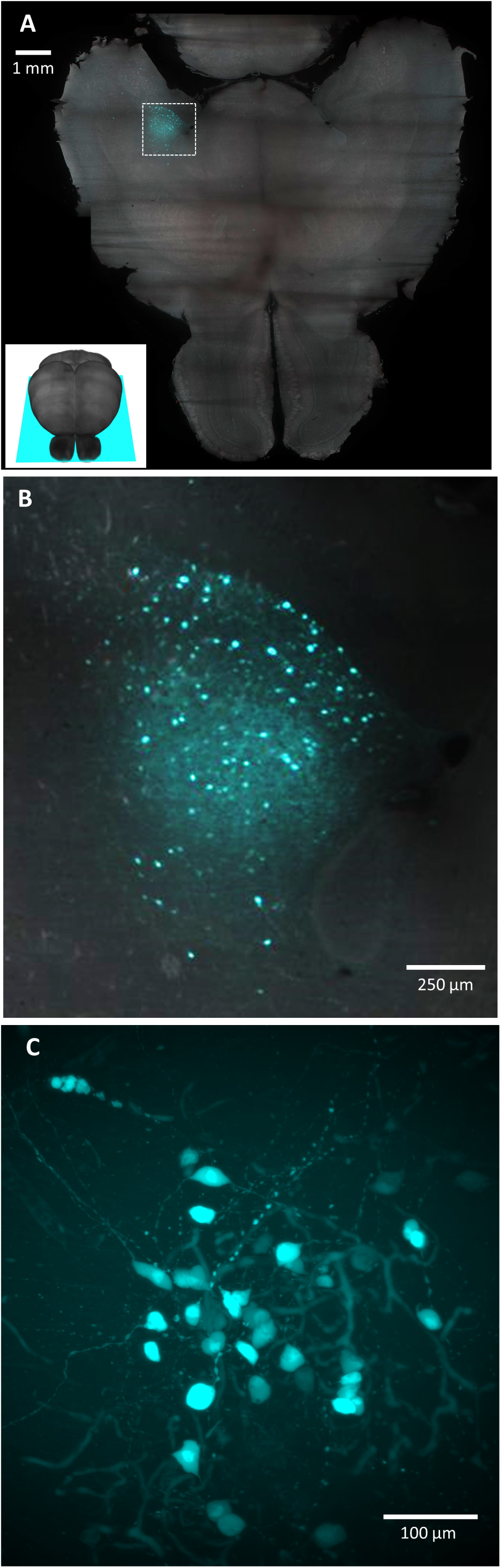
Light sheet microscopy with MHD-cleared tissue. A) Optical slice of an intact mouse brain, which was cleared using MHD-accelerated clearing in the horizontal orientation (inset: orientation of optical slice). GFP-expressing neurons are clear in the medial amygdala (cyan). B) A higher magnification image of the infection site corresponding to the location of the dashed box in the horizontal view from A). C) A higher magnification (20x) image of GFP-expressing neurons in an intact mouse brain imaged 3 mm from the ventral and lateral surface of the tissue in the bed nucleus of the stria terminalis.

**Figure 4:**
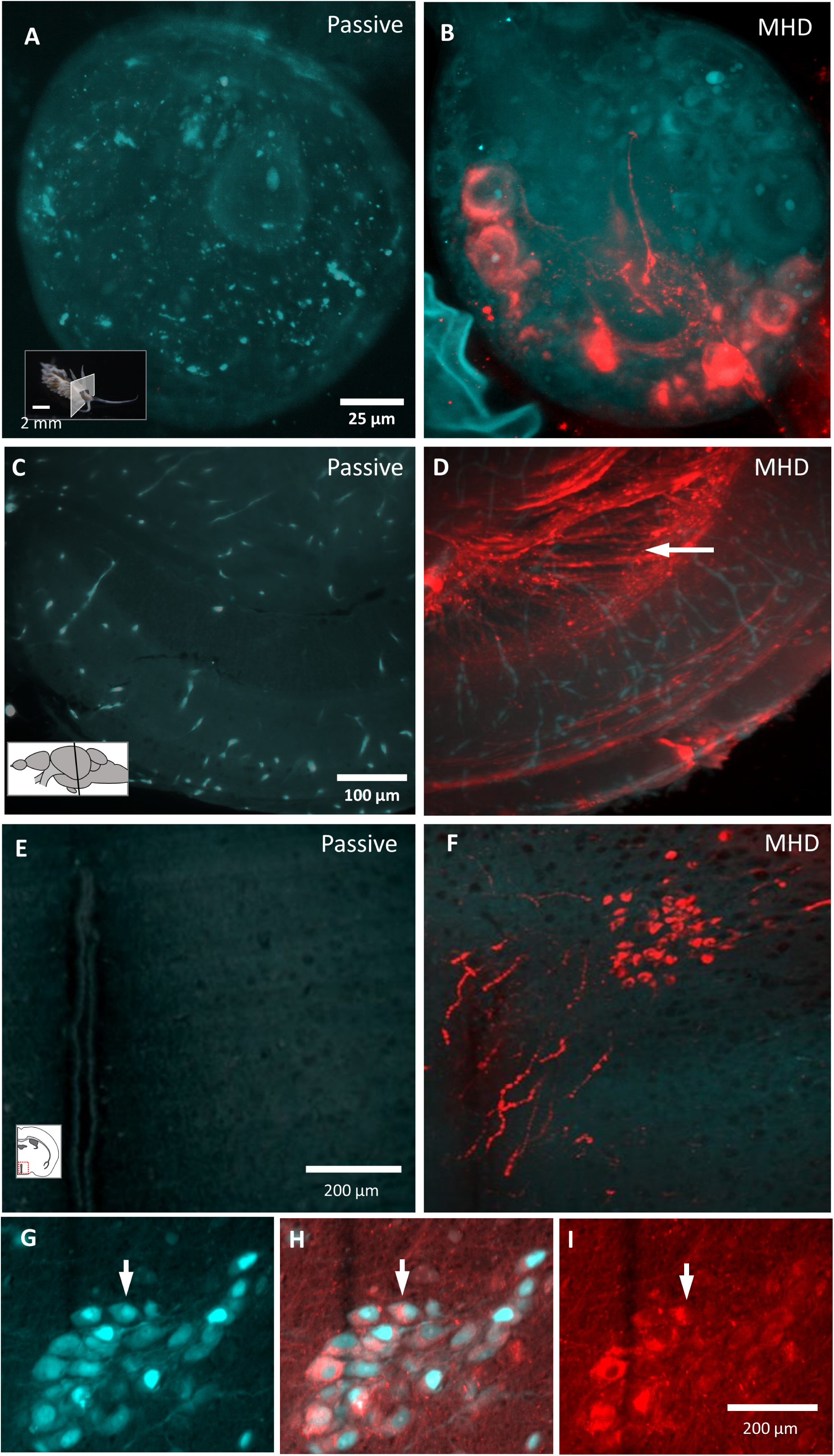
MHD-accelerated antibody labeling of brain tissue from sea slug, zebrafish, and mouse. A), B) 20x image of the pedal ganglion imaged inside the intact animal (Berghia stephanieae) after MHD-accelerated (B) and passive (A) α-serotonin antibody labeling (red) with tissue autofluorescence (cyan). C, D) Images of a cleared adult zebrafish brain (3mm × 3mm × 6mm) after α-acetylated tubulin antibody labeling (red) with tissue autofluorescence (cyan; Passive labeling: C; MHD-accelerated labeling: D). E, F) Images of cleared mouse brain sample (6mm × 6mm × 6mm) after α-oxytocin labeling (red) with tissue autofluorescence (cyan; Passive labeling: E; MHD-accelerated labeling: F). G,H,I) MHD-accelerated labeling of adult mouse brain sample (6mm × 6mm × 6mm) after α-vasopressin antibody labeling (cyan) with genetically encoded tdTomato in vasopressin-expressing neurons (red; AVP-cre X ai9). Insets indicate the imaging plane.

### Tissue Labeling

MHD-accelerated labeling dramatically improved antibody penetration and allowed labeling of large intact tissue samples (Figure 4 – Figure supplement 1, Figure 4 —Figure supplement 2). An intact adult nudibranch (*Berghia stephanieae*) (medio-lateral: 1.3 mm, dorso-ventral: 1.5 mm, anterio-posterior: 2 cm) that had been delipidated using the MHD-accelerated clearing device was incubated with an anti-serotonin (5-HT; Immunostar; 1:500) antibody followed by a fluorescent secondary (488 nm conjugated; ThermoFisher; 1:200). Passive incubation for 12 hours resulted in little to no penetration into the brain (Figure 4A), whereas MHD-accelerated antibody labeling for 12 hours drove antibodies throughout the sample and revealed 5-HT expressing cell bodies and projections (Figure 4B).

Intact zebrafish brains (medio-lateral: 3 mm; dorso-ventral: 3 mm; anterio-posterior: 6 mm) were passively delipidated in SDS for 7 days and then incubated with anti-acetylated tubulin antiserum (Immunostar; 1:500; Piperno, et al., 1985) for 12 hours to identify neural fibers (Figure 4 C,D). Control tissue samples (no MHD force applied) showed minimal antibody penetration along the outer edge of the tissue with little fluorescence visible in the optic tectum (Figure 4C). In contrast, MHD-accelerated labeling for the same amount of time showed robust labeling of neural tracts throughout the brain (Figure 4D).

To test MHD-accelerated labeling in mammalian tissue, an anti-oxytocin (OT) antibody was applied to a cube of mouse brain (medio-lateral: 6 mm, ventro-dorsal: 6 mm, antero-posterior: 6 mm) centered on the periventricular nucleus of the hypothalamus (1:500 primary; 1:200 secondary). As above, antibodies did not effectively penetrate the tissue sample in the absence of MHD force (Figure 4E). In contrast, OT-expressing cells were clearly visible in the PVN, located deep within of tissue cube, using MHD-accelerated labeling (Figure 4F). OT-expressing neuronal processes were easily resolved and were seen to project towards the third ventricle and, consistent with OT neuron morphology. Accurate OT-labeling was seen >1.8 mm from the nearest edge. The ability to visualize axonal varicosities and nuclei in OT-labeled neurons demonstrated that the MHD-accelerated labeling strategy can be used to resolve subcellular structures (Figure 4F).

To confirm the specificity of antibody binding is maintained in MHD-accelerated labeling, we used an anti-vasopressin antibody in mice that expressed tdTomato in vasopressin-expressing neurons (Figure 4 G-I). Tissue was generated by crossing the Ai9 Rosa26:LSL:tdTomato reporter line (Madisen, et al., 2010) and a line where Cre recombinase is expressed under the control of the arginine vasopressin (AVP) promoter (Bendesky, et al., 2017). This produced tissue where the fluorescent reporter tdTomato was expressed under the control of the AVP promoter. An example is shown for a 12-hour MHD accelerated antibody label. This produced specific co-labeling of the genetically encoded fluorophores and the anti-AVP antibody (Figure 4G-I).

## Discussion

New tissue clearing techniques, combined with fluorescent transgenic reporters and antibody labeling, allow unprecedented investigation of gene expression, neuronal connectivity, and functional anatomy in the brain. The MHD-accelerated protocol outlined here harnesses the strengths of hydrogel-based clearing approaches to maintain proteins and genetically encoded fluorescence in large samples (Chung, et al., 2013; Susaki, et al., 2015; Lee, et al., 2016; Pan, et al., 2016). MHD acceleration also uses electrophoresis to clear and label samples, but then adds to this an additional MHD force that produces coherent bulk flow of buffer and ions at the intersection of orthogonal magnetic and electrical fields (Qian, et al., 2009). Thus, the MHD force accelerates tissue clearing and antibody labeling without increasing the strength of potentially damaging electrical fields (Figure 4—Figure supplement 2). Additionally, MHD forces produce unidirectional buffer flow inside the tissue sample itself which helps regulate temperature in the center of the sample and to pull unbound molecules into and out of the sample.

The efficiency of MHD forces to rapidly drive charged molecules into and out of tissue is a consequence of a fundamental difference in the way that MHD fields and electrical fields act on charged particles. Electrophoresis drives cations and anions in opposite directions. In contrast, MHD-forces drive cations and anions in the same direction along the third orthogonal axis (Qian, et al., 2009). Because the MHD force acts on all ions is in the same direction, MHD creates a ‘flow’ in all charged molecules inside the field (Figure 1B). As a result, MHD forces generate rapid flow of buffer through the device and through a tissue sample located within the MHD field (Video 1). Indeed, the velocity produced by MHD increases linearly with voltage and far exceeds the nominal velocities produced in the same device when the magnets are removed (Figure 1). The induced flow of buffer dissipates local heating near the electrodes, and in the core of the tissue sample, that could otherwise damage tissue. Because MHD forces are cumulative to the maintained electrophoretic forces, the MHD approach described here maintains all the advantages of electrophoretic tissue clearing and labeling and then adds a complementary force that further propels lipids, antibodies, and buffer through the tissue.

MHD acts directly on ions at the intersection of electric and magnetic fields and both labeling and clearing devices described here exploit this feature. By placing the tissue directly at the intersection of these fields, a force is produced from inside the tissue (Figure 1C). The buffer flow induced by this force serves three purposes. First, in the case of tissue clearing it helps remove lipids from the tissue sample. Second, in the case of antibody labeling it helps push antibodies into the tissue. While MHD acts directly on antibodies, we believe that the strategy described here may be effective primarily because MHD generates something akin to a river of buffer flowing in the same direction through the microchannels in the hydrogel fixed tissue. Like twigs caught in the flow of a river, antibodies and lipids are pulled through the tissue sample allowing rapid clearing and labeling. Third, MHD driven buffer flow facilitates efficient thermal regulation of precious tissue samples. Instead of simply introducing cooler buffer solution to the surface of the tissue, the cooler solution actively replaces the existing warmer buffer within the tissue.This ensures that the center of the tissue will experience more similar conditions to the outer edges of the tissue than would be possible with only an electrical field. As a result, an endogenous pump is no longer required to maintain thermal equilibrium and a stronger force can be applied with less electrical current (Qian, et al., 2009). Moreover, because the ‘pumping’ action of MHD is produced directly from the electrical and magnetic fields without moving parts, it is virtually impossible for the pump to fail during the clearing or labeling process.

This approach eliminates the need for solvents that are harmful to fluorophores (e.g., methanol), and simplifies tissue clearing to the bare minimum components. Indeed, the only obligatory requirement is that the tissue sample is held at the intersection of an electrical and a magnetic field. Thus, the strategy outlined here is remarkably clean, efficient, and adaptable. The device itself can be 3D printed in plastic (Figure 5) making the device remarkably simple to build. Accordingly, with an investment of about $100 this device can be modified and specialized for the specific needs of a given experiment.

**Figure 5:**
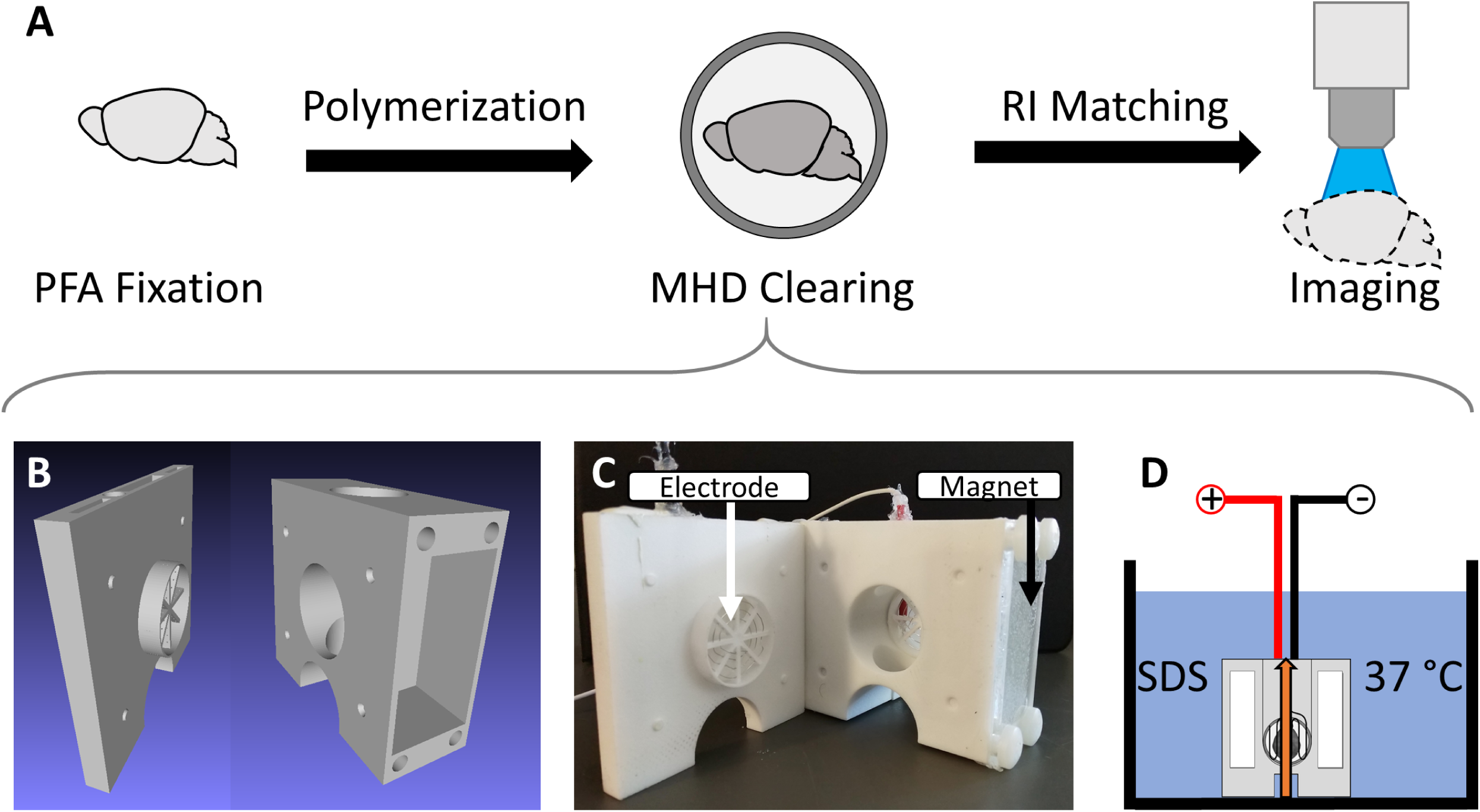
Overview of MHD-accelerated clearing approach. A) Steps required to effectively clear tissue of lipids. B) CAD diagram showing the MHD-assisted clearing device. C) A photograph of the clearing device with tissue chamber exposed and arrows to show the location of the magnets and electrodes in the device. D) A cartoon showing the setup of the clearing device submerged in a container filled with detergent solution held at 37 °C. Tissue was placed in the central chamber where MHD force (orange arrow) produced from the electrical and magnetic fields simultaneously circulate the buffer solution and accelerate clearing.

The MHD-based approaches described here will reliably allow rapid tissue clearing and antibody labeling of large intact tissue samples, rendering them suitable for 3D fluorescent imaging. We demonstrate the efficacy of a simple MHD device by clearing and labeling sea slug, zebrafish, and mouse tissue and using multiple antibodies. Each of the antibody labeling procedures used here required no more than 4.5 mL (1:200 concentration) of labeling solution which can be collected at the end of the procedure and reused. Combined with the linear rate of antibody penetration observed with longer durations of active labeling, we believe this system can be adapted for the fluorescent labeling of even larger samples, with the limit ultimately constrained by the imaging system employed and not by the ability to produce high-quality tissue samples.

## Methods

### Animal Use

All vertebrate animals were handled according to a protocol approved by the UMass Amherst Institutional Animal Care and Use Committee (IACUC; protocol #2018-0014 and #2017-0060).

### Measure of MHD-induced flow

A solution of sodium chloride was made in a small tank (2.5 L). Sodium chloride was slowly added to the tank until the electric conductivity of the solution matched that of the clearing solution. The clearing device was then submerged in the solution with a measured grid behind the tank to provide scale. 0V, 10V, 20V, 30V, 40V, 50V, or 60V were then applied to the device and sodium alginate spheres were introduced into the tank at a constant location (N = 7). The velocity of the spheres through the device was measured. Velocity was calculated using a high-speed video taken over a calibrated grid. This process was then repeated using only an electric field (without magnets). Paired-sample t-tests were performed between the MHD and electric-only conditions at each voltage using MATLAB. The p-values were corrected for multiple comparisons using Bonferroni correction. Each condition was fit to a linear model using MATLAB.

### Design of MHD-accelerated clearing device

The strategy for using MHD to remove lipids from tissue samples requires binding proteins and polymerizing a hydrogel, removing lipids, and matching the refractive index of the tissue and imaging media (Figure 5A). A tissue chamber was placed into the central chamber of the MHD-accelerated clearing device (Figure 5B, C). This holds the tissue at the intersection of the electrical and magnetic fields. The clearing chamber was submerged in a large (5 L) bath of clearing solution at 37 °C and 30 VDC (0.25 Amps) was applied across the tissue for several hours (typically 16 hours for mouse brain tissue and 2 hours for intact zebrafish brains; Figure 5D).

### Tissue Fixation and hydrogel polymerization

Mice were anesthetized with isoflurane, euthanized, and perfused with 0.01 M phosphate buffered saline (PBS) followed by 4% paraformaldehyde (PFA) in 0.01M PBS. Tissue was then post-fixed in 4% PFA at 4 °C overnight. Next, the tissue was placed in a hydrogel solution (4% acrylamide, 4% PFA, 0.05% bis acrylamide, and 0.25% VA-044 initiator suspended in 0.01 M PBS) at 4 °C overnight (Chung, et al., 2013; Isogai, et al., 2014). Oxygen was flushed out of hydrogel-infused tissues nitrogen gas and then the samples were polymerized by incubating them at 37 °C overnight (Chung, et al., 2013). Excess hydrogel was removed from the surface and tissue samples were transferred to PBS to flush hydrogel monomers.

Adult zebrafish were euthanized in 0.2 mg/ml tricaine mesylate (MS-222), decapitated, and the heads placed in 4% paraformaldehyde overnight. Heads were then placed in PBS and brains were carefully dissected, incubated in hydrogel at 4 °C overnight, and processed as above.

Adult nudibranchs (*Berghia stephanieae*) were anaesthetized in cold 4.5% magnesium chloride in artificial sea water for 20 minutes, pinned to a Sylgard-lined dish, and fixed in 4% paraformaldehyde in sea water overnight at 4 °C. Whole animals were washed with PBS and then incubated in hydrogel at 4 °C overnight and processed as above.

### Active Tissue Delipidation (clearing)

Tissue samples were incubated in SDS-clearing solution (10 mM sodium dodecyl sulfate in 0.1 M borate buffer, pH 8.5) for 2 days at 37 °C unless otherwise noted. Samples were then transferred to the MHD-accelerated clearing chamber, consisting of two interlocking cell-strainers (ThermoFisher; catalog #: 87791). This chamber was placed in the intersection of the electrical and magnetic fields in the center of the device (Figure 5) and the chamber was lowered into a bath of 37 °C SDS. 30V DC were then applied across the tissue to initiate MHD-accelerated clearing. After clearing, the tissue is taken out of the clearing chamber and washed in 0.1 M PBS for at least 12 hours.

### Refractive Index Matching and Light Sheet Microscopy

The tissue was transferred to “Optiview” (Isogai, et al., 2014) refractive index matching solution and incubated at 37 °C for at least 12 hours to achieve optical clarity through RI matching (Figure 5A; Isogai, et al., 2014). Samples were imaged at 5X or 20X magnification with a lightsheet microscope adapted for a 1.45 RI imaging solution (Zeiss Z1).

### Measures of Clearing Efficacy

36 mouse brains were embedded in hydrogel, cleared using the MHD-accelerated clearing protocol, and assessed for transparency. The tissue was divided into two groups: one that was pretreated by passively delipidating in SDS clearing solution for two days at 37 °C, and a second that was placed in a 0.1 M borate buffered solution at 37 °C for the same time as the pretreatment. Tissue samples from each condition (n=3) were then actively delipidated using the MHD-accelerated clearing system for 2, 6, 8, 12, or 15 hours. After washing in 0.01M PBS, the tissue was equilibrated in Optiview (Isogai, et al., 2014) for 48 hours at 37 °C.

Transparency was determined by the percentage of light transmitted through the tissue and the maximum depth from the external surface at which the morphology of neural processes (including primary dendrites and axons) could be resolved. Light transmission was measured using a wide-spectrum light-source and calibrated photodiode. Data from each condition was fit with a saturating exponential curve in MATLAB.

### MHD-accelerated staining of fixed tissue with methylene blue

Penetration of methylene blue into a 1 cm^3^ cube of homogeneous brain tissue under MHD force was tested over 1, 2, and 4 hours (N = 1). Cubes of uncleared sheep brain tissue were equilibrated to the antibody labeling buffer solution for 12 hours. The tissue was then placed at the intersection of a strong magnetic and electric field (30V DC) and submerged in a solution of methylene blue (0.1 M) buffered to pH 9.5 (37 °C). The orientation of the electric field was reversed at 15-minute intervals for 3 minutes. Three samples were labeled using this approach for 1, 2 or 4 hours. Following the stain, the tissue was bisected and imaged. A control sample was incubated in the same solution (37 °C) for 4 hours without the application of any active force. This sample was bisected and imaged as the others.

### Comparative staining of methylene blue into agarose cubes as a result of various strengths of electrical force conjugated to MHD force

15 1 cm^3^ of 3 % agarose were subjected to labeling methylene blue labeling by MHD force for 0, 5, 10, 15, 30, 60, or 120 minutes at varied electrical field strengths. The distance penetrated into the agarose cubes was measure after bisection and plotted against staining time with 10, 20, or 30V in a constant magnetic field.

### Optimization of MHD-Accelerated Immunohistochemistry

0.22 cm^3^ cubes of hydrogel were incubated with FITC-conjugated antibodies (1:200; Jackson Immunoresearch) in a buffered solution (pH 9.5) with MHD assistance, with an equivalent electrical field (30V DC), or passively for 1 hour at 37 °C (N = 1). These cubes were incubated at the center of the intersection of the electrical and magnetic fields or the center of the electrical field. After labeling samples were RI-matched using “Optiview” (Isogai, et al., 2017) and imaged using a Zeiss Z1 light-sheet microscope at 5x magnification. The farthest distance from the edge of the tissue in the orientation of the MHD and electrical force where fluorescent antibody was observed determined the penetration of the antibody.

### Antibody Labeling

Delipidated tissue was placed inside of a 2-inch length of 0.25-inch diameter dialysis tubing (6 – 8 kDa); Spectrum). After equilibration, samples were incubated in an antibody solution inside dialysis tubing at the center of intersecting electrical and magnetic fields where the MHD force was strongest (Figure 6). Confining the tissue sample inside dialysis tubing reduced the volume of antibody required for labeling and protected the tissue sample and antibody solution from direct exposure to the electrodes. Magnets (Applied magnets; NB057-6-N52) were placed on the top and bottom of the MHD labeling device creating a central chamber Figure 6B). The ends of the dialysis tubing were connected to 9.5 mm diameter vinyl tubing (ThermoFisher: S504591) using 0.25-inch Leur lock barbs (Cole-Parmer; UX-45501-20) to create a torus-shaped chamber allowing the antibody solution to circulate continuously and provide an even and continuous source of antibody to the tissue sample (Figure 6). Antibody solution (4.5 mL; 0.1 M borate buffer titrated to pH 9.5 with 0.1 M LiOH, 1% heparin, 0.1% Triton X-100; 1:500 primary antibody) was transferred into the dialysis tubing using a 5 mL syringe. The labeling chamber was submerged in a 1L tub containing 0.1M Borate Buffer pH 9.5/0.1% Triton X-100 solution. A 5 mL syringe filled with the buffer solution was attached to the circulation line to maintain constant pressure inside of the dialysis tube. 60 volts DC (∼0.2 Amps) was applied across the electrodes for 15 minutes, followed by 3 minutes of inactivity repeatedly for 12 hours to drive antibodies into the tissue sample. The system was held at 37 °C and protected from ambient light to minimize bleaching of fluorophores throughout the procedure.

**Figure 6:**
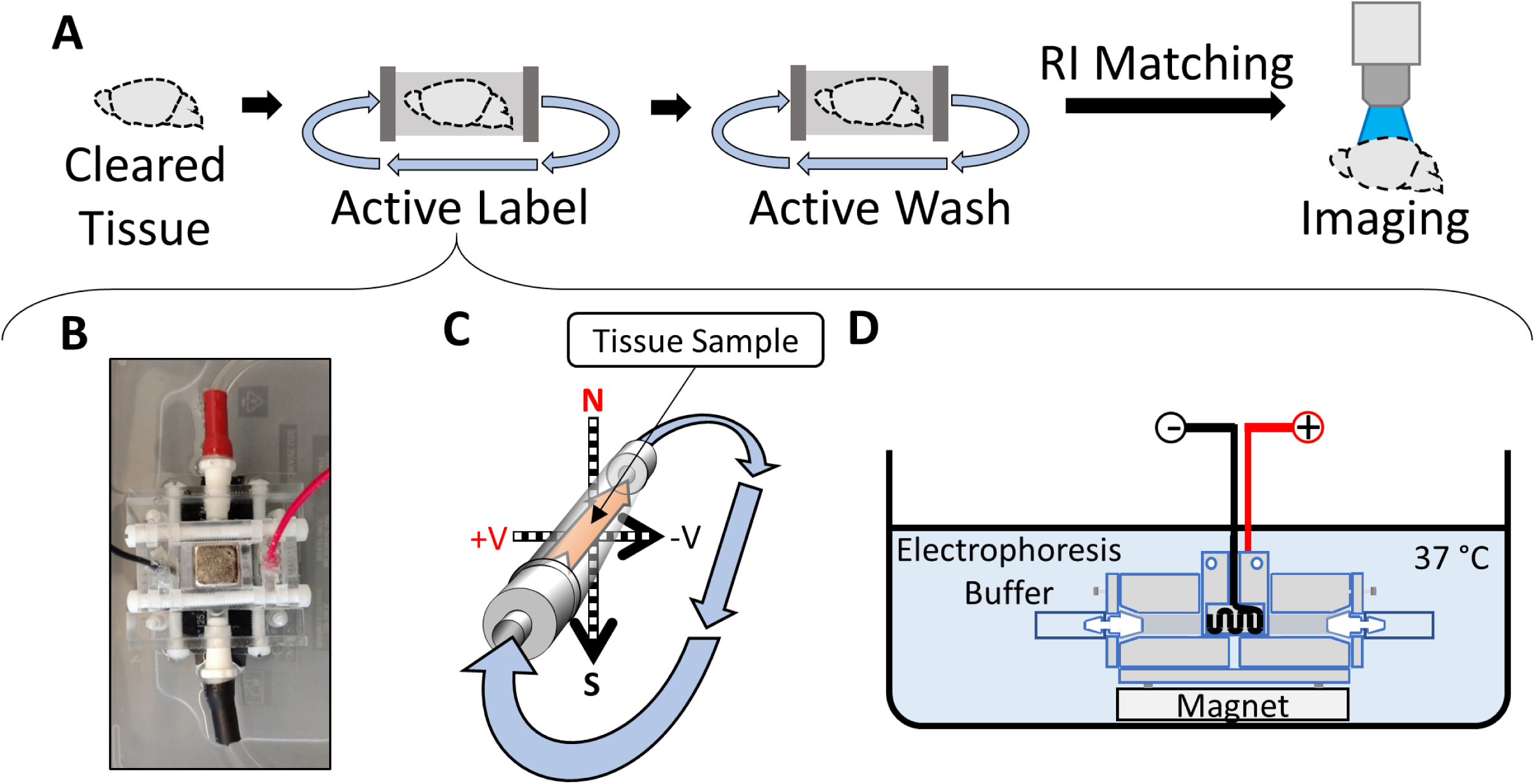
Overview of MHD-accelerated antibody labeling. A) Illustration of the steps required to label and image tissue. B) Picture of the MHD-assisted labeling device. C) Schematic showing the tissue location inside the MHD-assisted labeling device. The direction of the MHD force is indicated by the orange arrow inside the dialysis tubing. The resulting direction of flow of the solution through the closed loop is indicated by the blue arrows. D) Diagram of the antibody labeling device setup for a label. The device is submerged in a bath of electrophoresis buffer held at 37 °C.

Following each round of MHD-accelerated labeling, the antibody solution was replaced with a wash solution (0.1 M borate buffer titrated to pH 9.5 with 0.1 M LiOH, 1% heparin, 0.1% Triton X-100) and the tissue was exposed to 6-hours of “active washing” using the same voltage settings. Labeled tissue was then washed in 0.01 M PBS for at least 12 hours.

## Acknowledgements

We thank D. Kelly, P. Sterling, and the members of the Bergan lab for helpful comments on this paper. Support for these experiments came from the University of Massachusetts at Amherst, the Armstrong Fund for Science (J.F.B), and a generous gift from Britton Sanderford (J.F.B). J.F.D is a recipient of the University of Massachusetts at Amherst NSB Fellowship.

---

**Figure 2 —Figure supplement 1:**
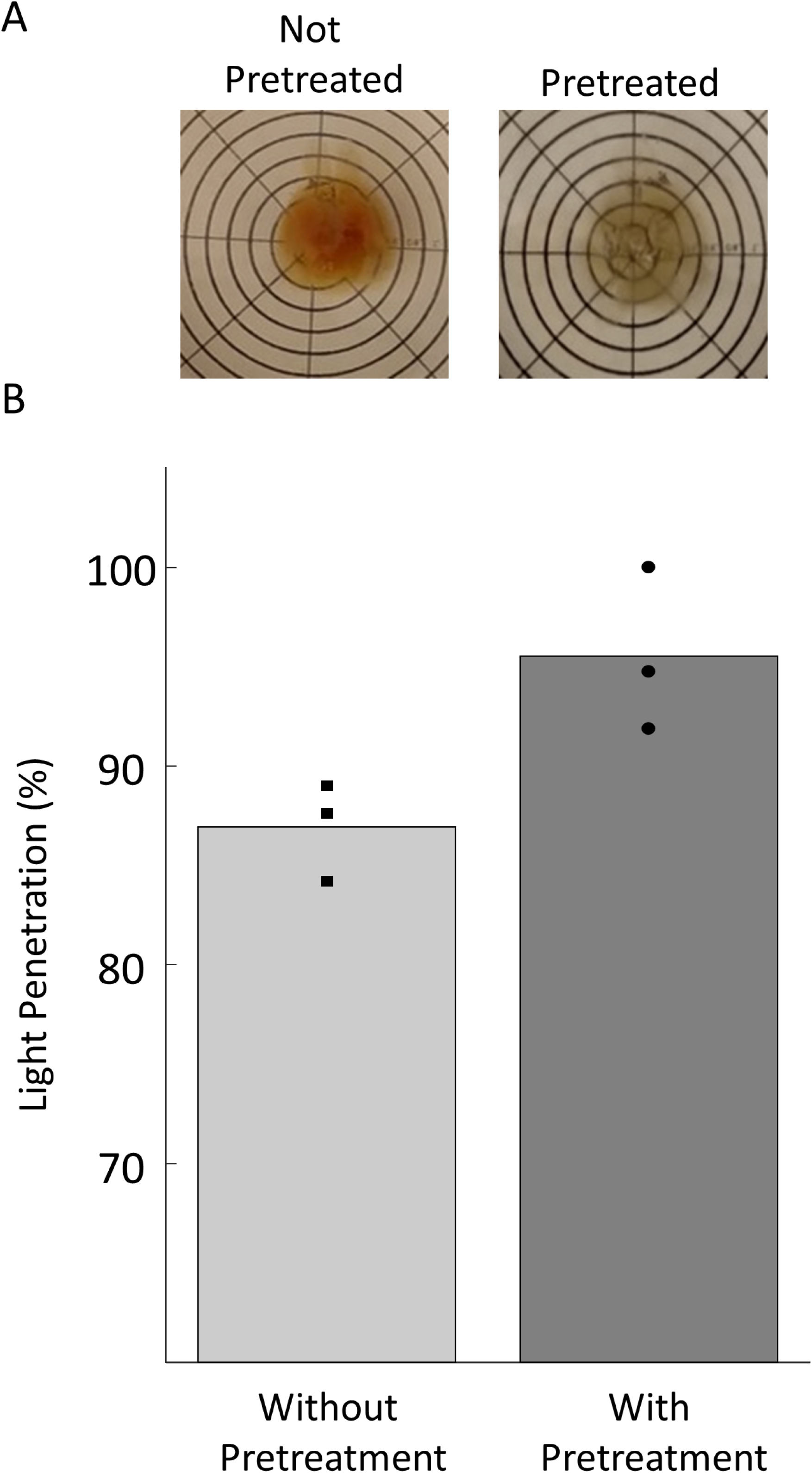
Comparison of tissue cleared with passive pretreatment and without pretreatment. A) Examples of brains cleared with MHD-accelerated clearing for 12 hours and then equilibrated in RI-matching solution with (right) and without (left) passive treatment in the clearing solution for 48 hours (37 °C) prior to MHD-accelerated clearing. B) Measurements of the optical transparency of brains that were actively cleared for 12 hours with and without passive treatment (N = 3). Transparency was measured as percentage wide-spectrum light penetration through the tissue (curve fit with a saturating exponential).

**Figure 4 —Figure supplement 1:**
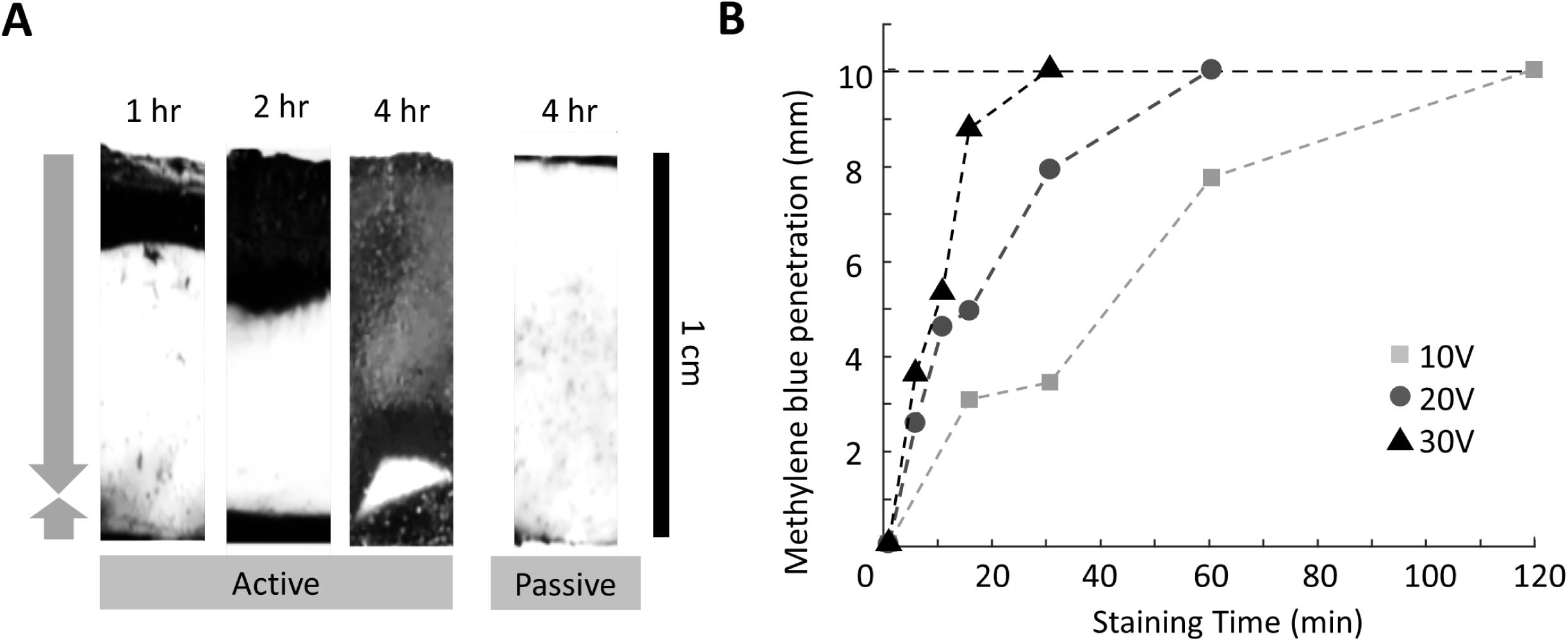
A) Penetration of methylene blue into a 1 cm^3^ cube of homogeneous brain tissue as a result of MHD force over 1, 2, and 4 hours (N = 1). The fourth image shows a comparative 4-hour stain without MHD force. The arrows on the left-hand side of the images demonstrate the direction of the MHD force with respect to the tissue. The length of the arrows demonstrates the proportion of time when the MHD force was aimed in the direction indicated by each arrow. B) shows the comparative staining of methylene blue into agarose cubes as a result of various strengths of electrical force conjugated to MHD force. The distance the methylene blue penetrated into the agarose cubes is measured against the amount of time stained with 10, 20, or 30V conjugated to a constant magnetic field. Note: panels A and B are from different iterations of the staining device and the rate of penetration was significantly improved for panel B.

**Figure 4 —Figure supplement 2:**
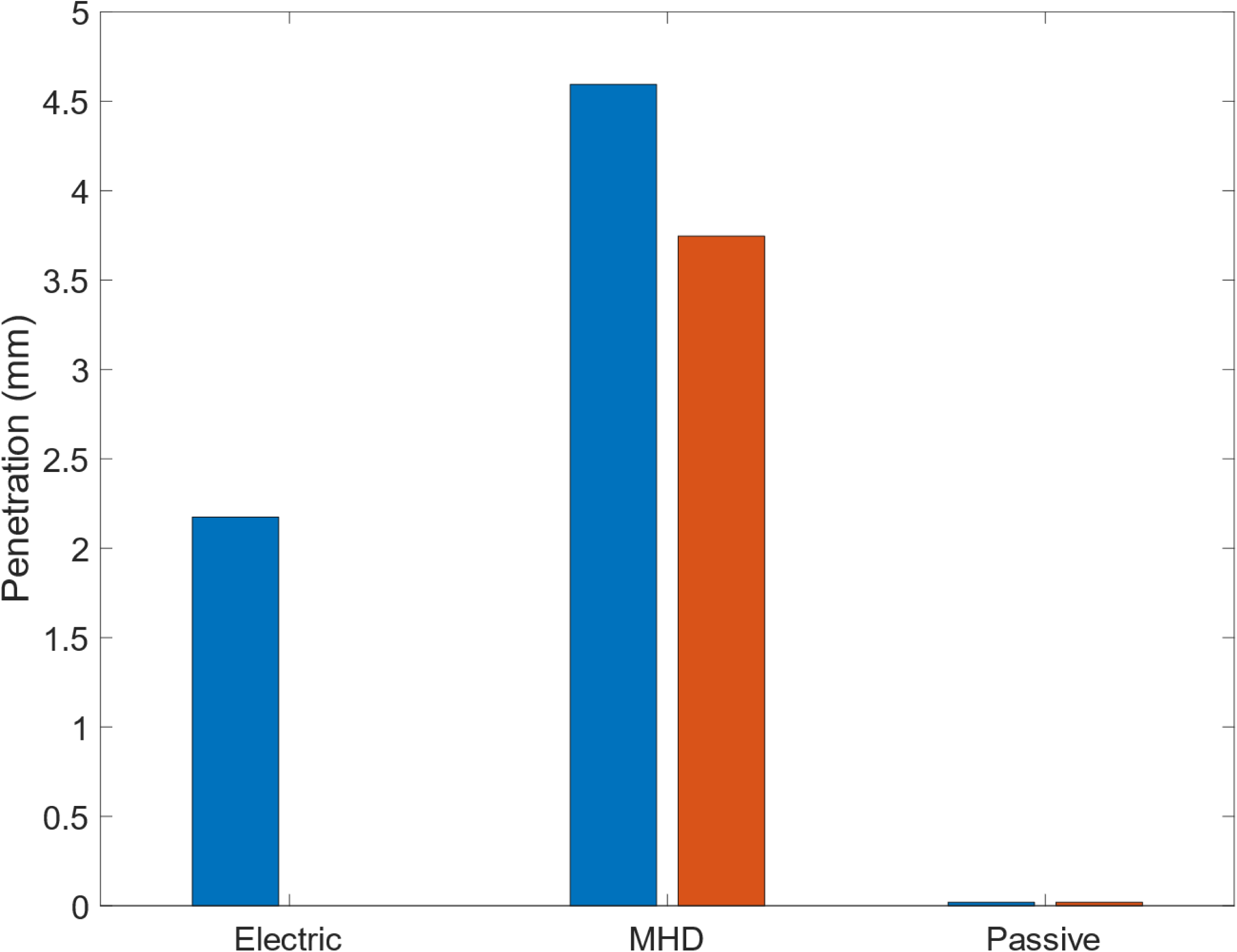
The distance of antibody penetration from the nearest surface after 1 hour of MHD-accelerated labeling, electrophoretic labeling, or passive labeling (N = 1) was measured using light-sheet microscopy. The blue and orange bars represent the distance the antibody penetrated the sample in the orientation of the electric (blue) or MHD (orange) force.

